# Bridging 3D Molecular Structures and Artificial Intelligence by a Conformation Description Language

**DOI:** 10.1101/2025.05.07.652440

**Authors:** Jiacheng Xiong, Yuqi Shi, Wei Zhang, Runze Zhang, Zhiyi Chen, Chuanlong Zeng, Xun Jiang, Duanhua Cao, Zhaoping Xiong, Mingyue Zheng

**Author notes:** Correspondence author. Email address (Mingyue Zheng). These authors contributed equally to this work (Jiacheng Xiong, Yuqi Shi).

## Abstract

Artificial intelligence, particularly language models (LMs), is reshaping research paradigms across scientific domains. In the fields of chemistry and pharmacy, chemical language models (CLMs) have achieved remarkable success in two-dimensional (2D) molecular modeling tasks by leveraging one-dimensional (1D) representations of molecules, such as SMILES and SELFIES. However, extending these successes to three-dimensional (3D) molecular modeling remains a significant challenge, largely due to the absence of effective 1D representations for capturing 3D molecular structures. To address this gap, we introduce ConfSeq, a novel molecular conformation description language that integrates SMILES with internal coordinates including dihedral angles, bond angles, and pseudo-chirality. This design naturally ensures SE(3) invariance, while preserving the human readability and conciseness characteristic of SMILES. ConfSeq enables the reformulation of a range of 3D molecular modeling tasks, such as molecular conformation prediction, 3D molecular generation, and 3D molecular representation, into sequence modeling problems. Then, by simply employing a standard Transformer architecture, we achieve state-of-the-art performance on various benchmark sets. Furthermore, compared to widely used diffusion-based approaches in 3D molecular modeling, the ConfSeq-based method offers unique advantages in inference efficiency, generation controllability, and enables scoring of generated molecules. We believe that ConfSeq can serve as a foundational tool, advancing the development of sequence-based 3D molecular modeling methods.

## Main

Artificial intelligence (AI) is profoundly transforming research paradigms across various fields^1^. Among these, language models (LMs), characterized by their powerful generative capabilities, broad adaptability, and human interaction capabilities, have emerged as the most influential and disruptive AI technologies^2-5^. In recent years, LMs have been widely applied in the development of general large models, like GPT-4^6^ and Gemini^7^, as well as domain-specific large models, like ESM^8,9^ and GeneFormer^10^. This has significantly advanced scientific research, industrial applications, and social transformation.

In pharmacy and chemistry, chemical language models (CLMs) have also become a key research focus and have been widely applied^11,12^. By encoding molecular topological structures as one-dimensional (1D) sequences (such as SMILES and SELFIES), CLMs can tackle nearly all two-dimensional (2D) molecular modeling tasks, including molecular translation^13,14^, molecular generation^15^, and molecular representation^16^, while delivering state-of-the-art performance across these tasks. However, molecular properties are determined not only by two-dimensional (2D) topology but also by three-dimensional (3D) geometry, which is essential for understanding and predicting molecular physicochemical properties^17^, chemical reactivity^18^, and binding modes and affinities toward biological targets^19^. Therefore, developing effective methods for the representation and generation of 3D molecular structures is essential for advancing AI-driven applications in chemistry and the life sciences. However, unlike the widespread success in 2D molecular modeling, 3D molecular modeling tasks have long been a restricted domain for CLMs, primarily dominated by various graph-based diffusion models^20-23^. Yet, these models often encounter significant challenges, including complex model architectures, low inference efficiency and controllability, and a lack of capabilities for ranking and scoring results. Therefore, further exploring the application of CLMs in 3D molecular modeling is both necessary and urgent.

In 2024, StoneWise introduced the Lingo3DMol model, which leverages LMs to generate SMILES of molecular fragments and incorporates additional modules to predict their spatial positions, ultimately constructing 3D molecules^24^. While effective to some extent, this reliance on auxiliary spatial modules results in an architecture incompatible with other LMs, limiting its generalizability and scalability. Moreover, this approach neglects the conformational flexibility inherent to molecular fragments, potentially compromising its accuracy in modeling 3D structures. More recently, some researchers have directly encoded the 3D coordinates of molecules as strings and concatenated them with SMILES, achieving 3D molecular generation based purely on LMs^25-28^. However, this approach overlooks the SE(3)-invariance of molecules, which could undermine the model’s stability. Additionally, given the limited ability of language models to process complex numerical data^29,30^, directly encoding 3D coordinates as strings, may hinder their ability to capture spatial positions and geometric relationships. Furthermore, these methods often lack thorough comparisons with alternative approaches across diverse benchmark datasets, failing to fully elucidate the advantages of language models in 3D molecular modeling.

We argue that a key challenge in adapting language models for 3D molecular modeling is the absence of an ideal language for representing molecular conformations. Such a language should be capable of discretizing and sequentializing continuous, unordered geometric information while maintaining SE(3) invariance. Furthermore, for convenience and efficiency of use, it should also remain concise and human-readable. To this end, we introduce ConfSeq, a novel conformation description language. It encodes molecular conformations using some key internal coordinates including torsion angles, bond angles, and our newly proposed pseudo-chirality, which inherently preserves SE(3) invariance. These angular values lie within the closed interval of –180 to 180 degrees and can be efficiently discretized into a finite set of tokens. These tokens are ordered according to the atom and bond sequence in SMILES and are embedded at corresponding positions to generate ConfSeq. This design renders ConfSeq highly readable and facilitates data augmentation techniques analogous to SMILES enumeration (Fig. 1).

**Fig. 1.**
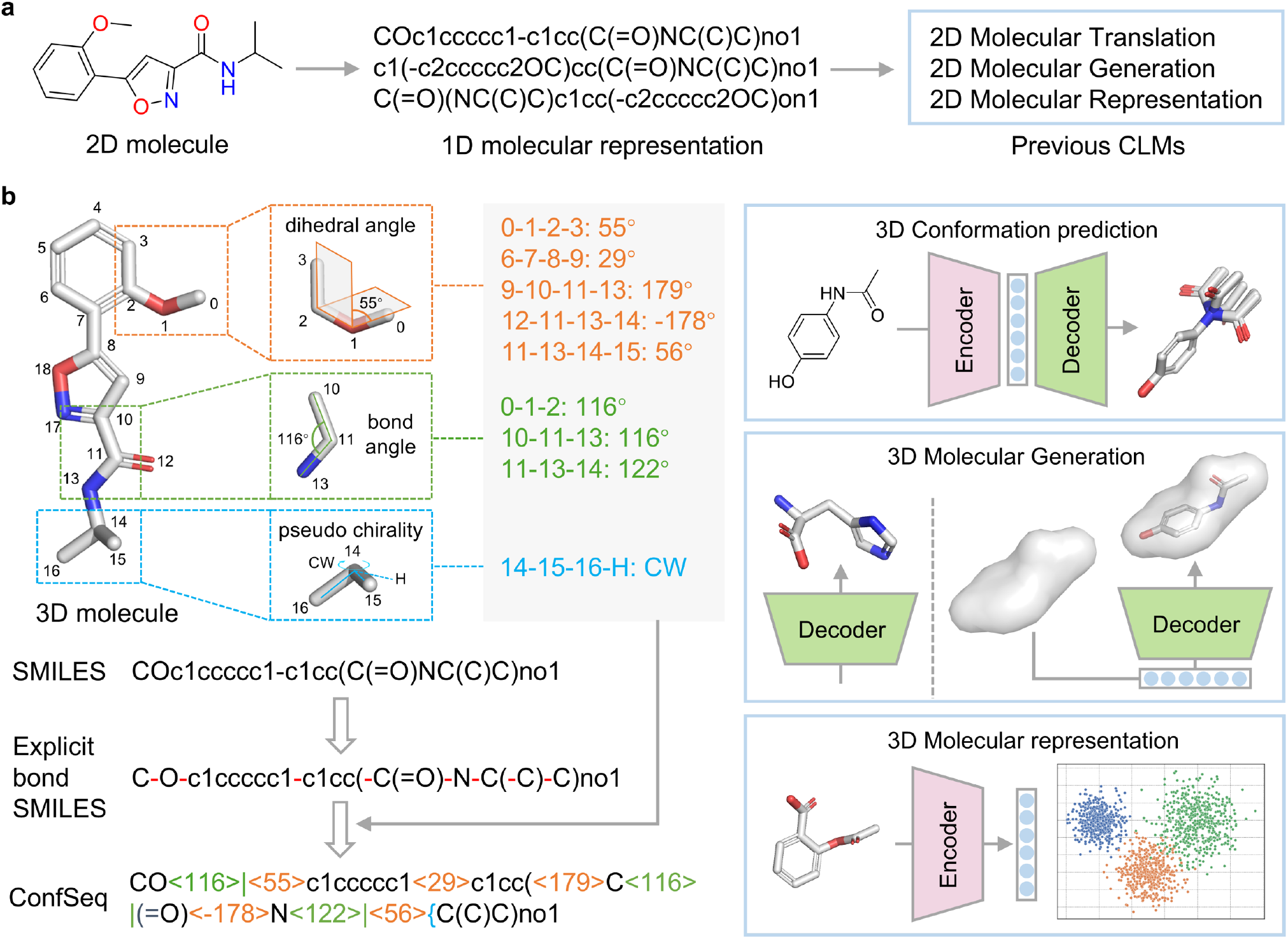
Overview of this work. (a) Previous work on 2D molecular modeling using chemical language models. (b) 3D molecular modeling using our proposed molecular conformation description language.

Using ConfSeq, we successfully reformulate a variety of 3D molecular modeling tasks, including 3D conformation prediction, 3D molecular generation, and 3D molecular representation, into sequence modeling problems. By employing a standard Transformer model, we achieve state-of-the-art performance across these tasks, significantly outperforming other elaborate graph-based approaches. Moreover, this sequence-based approach to 3D molecular modeling exhibits several unique advantages over existing methods, including higher computational efficiency, greater flexibility and controllability in the generation process, and the ability to score and rank generated candidates. These results collectively demonstrate the practicality of ConfSeq and highlight the potential of language models as a promising new paradigm for 3D molecular modeling.

## Results and Discussion

### Molecular 3D Conformation Prediction with ConfSeq

To demonstrate the applicability of ReactSeq, we first applied it to molecular conformation prediction. A standard Transformer was employed to construct our conformation prediction model, which translates molecular SMILES into ConfSeq representations. To further enhance model performance, we adopted data augmentation and test-time augmentation techniques proposed in previous studies. The model was trained and evaluated on the GEOM-Drugs dataset, with Coverage (COV) score at the 0.75 Å threshold, and Matching (MAT) scores used to assess its Precision (P) and Recall (R).

As shown in Fig. 2a and 2b, our model outperforms all baseline methods across all evaluation metrics, with particularly substantial gains in Precision. Specifically, compared to the previous state-of-the-art method (Tor. Diff.)^20^, our model improves COV-P from 47.9% to 58.4% and reduces MAT-P from 0.86 Å to 0.77 Å (Supplementary Table 1). Since the number of rotatable bonds is a key factor influencing the difficulty of molecular conformation prediction, we further analyzed performance across molecules with different numbers of rotatable bonds. As shown in Supplementary Fig. 1, our model consistently achieves the best performance across the vast majority of rotatable bond ranges. We also further analyzed our model’s Coverage at different thresholds, as shown in Supplementary Fig. 2. Our model consistently achieves the highest COV-R and COV-P scores at thresholds below 1 Å and 1.5 Å, respectively. These results demonstrate the robustness of our model.

**Fig. 2.**
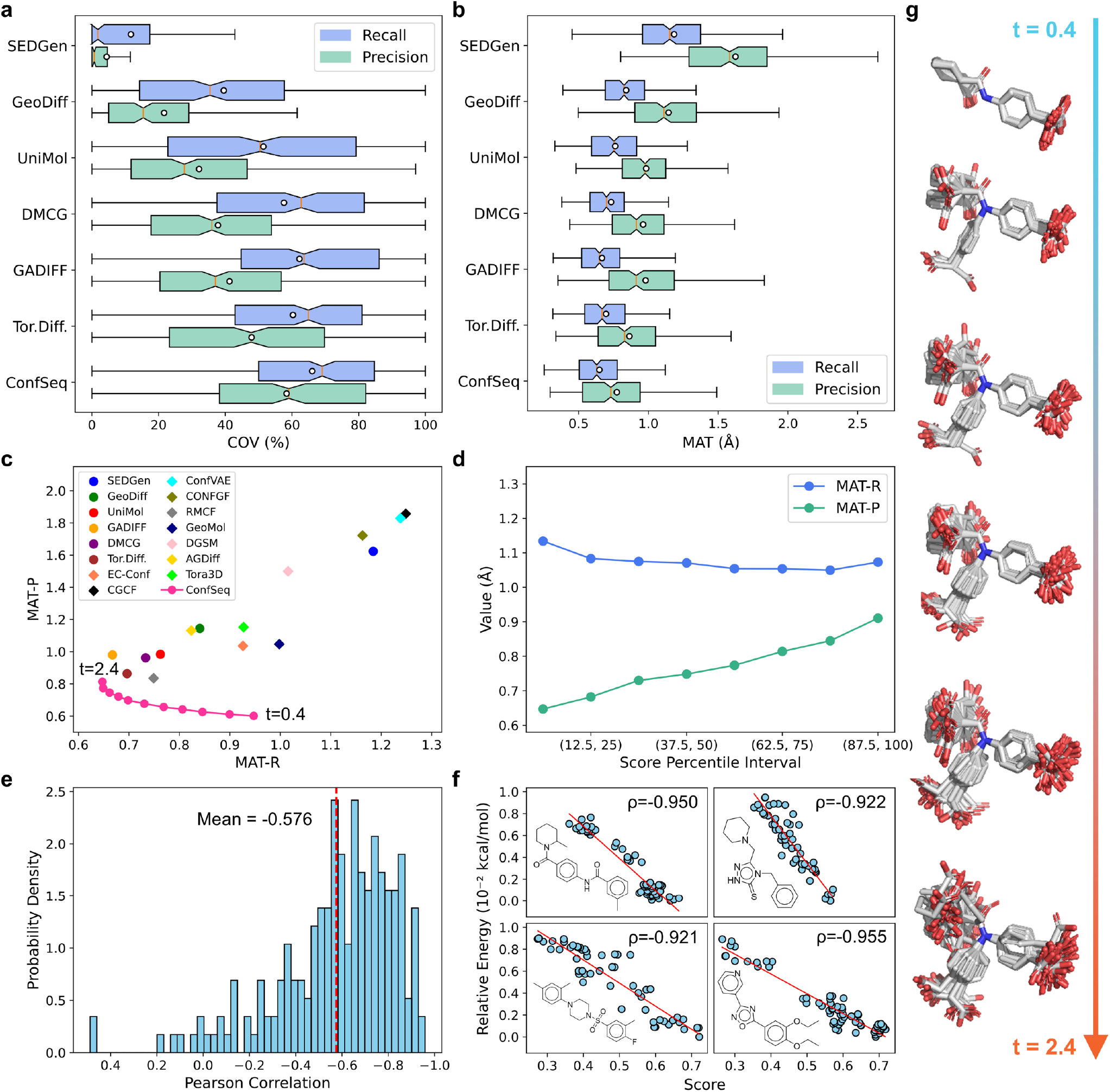
Performance of ConfSeq in molecular conformation prediction. (a-b) Comparison of Coverage and Matching scores between ConfSeq and baseline models. Circles in the box plots represent the mean values. (c) MAT-P and MAT-R scores of ConfSeq and other models across different sampling temperatures. Circles represent results from our experiments, while diamonds indicate values reported in previous studies. (d) MAT-R and MAT-P scores of generated conformations with different confidence scores. (e) Distribution of Pearson correlation coefficients between model-assigned confidence scores and conformational energies calculated by DFT. The red line indicates the mean value. (f) Representative molecules showing the strongest negative correlation between confidence scores and DFT-calculated conformational energies. “ρ” denotes the Pearson correlation coefficient. (g) Representative conformations generated by ConfSeq under different sampling temperatures.

Similar to other natural language models, our approach enables control over the diversity of predicted outcomes by adjusting the sampling temperature during generation. As shown in Fig. 2c, higher temperatures lead to improved Recall but reduced Precision, indicating a clear trade-off. The MAT-P and MAT-R scores of our model across different temperatures trace a curve consistently lying to the lower left of those from other models. This suggests that, in comparison to other models, our approach provides greater flexibility, allowing users to adjust the balance between Recall and Precision according to specific application requirements. Moreover, our model consistently delivers optimal performance, regardless of preference for precision or coverage. An example of conformations generated by our model at different temperatures is shown in Fig. 2g.

Beyond controllability, another distinctive advantage of our model lies in its ability to assign quantitative confidence scores to predicted conformations based on the stepwise probabilities computed during the autoregressive generation process. We first analyzed the confidence scores of molecules generated by the model in the conformation prediction task. As shown in Fig. 2d, as the confidence score increases, the MAT-R of the generated conformations remains stable, while MAT-P decreases substantially. This trend indicates that the model is capable of selecting higher-quality generated conformations based on its own scoring mechanism. Additionally, we further evaluated the model’s scoring capability on DFT-optimized conformations from the test set. As shown in Fig. 2e, the model’s confidence scores exhibit a strong negative correlation with molecular energy for the majority of molecules, achieving an average Pearson correlation coefficient of -0.58. Molecules with fewer rotatable bonds tend to show lower correlation coefficients (Supplementary Fig. 3), and for some, the correlation exceeds -0.9 (Fig. 2f). These results suggest that, through self-supervised learning on conformation data, our model has exhibited emergent intelligence in the understanding of the relationship between molecular geometry and energy, effectively capturing the underlying conformational energy landscape of molecules.

### Unconditional 3D molecular generation with ConfSeq

Molecular generation plays a pivotal role in de novo drug design, enabling efficient exploration of the vast chemical space beyond traditional trial-and-error approaches. We first evaluated ConfSeq on an unconditional 3D molecular generation task, which aims to learn the distribution of the training data and generate valid 3D molecular structures from scratch. A decoder-only Transformer was employed to construct our model, which was compared against state-of-the-art diffusion-based methods.

As shown in Table 1, with respect to 2D evaluation metrics such as validity, uniqueness, and their product, our model achieves nearly 100% chemical validity while maintaining high uniqueness, significantly outperforming existing counterparts. To assess the quality of the generated three-dimensional structures, we employed two evaluation metrics. The first is PoseBuster validity (PB-validity), originally proposed in the context of molecular docking, which evaluates the geometric and energetic plausibility of molecular conformations^31^. The second is the minimal root-mean-square deviation (RMSD) between the generated conformations and a set of 100 reference conformers predicted using the RDKit toolkit with the MMFF94 force field, reflecting the structural deviation between the generated and force field-optimized conformations. Our method achieved a PB-validity of 0.83, representing a 0.06 improvement over the previous optimal approach. Additionally, our model achieved a minimal RMSD of 0.1024, which is comparable to that of the training set and 0.05 lower than the previous state-of-the-art methods. These results collectively indicate that our method generates three-dimensional molecular structures with superior conformational plausibility.

**Table 1.**
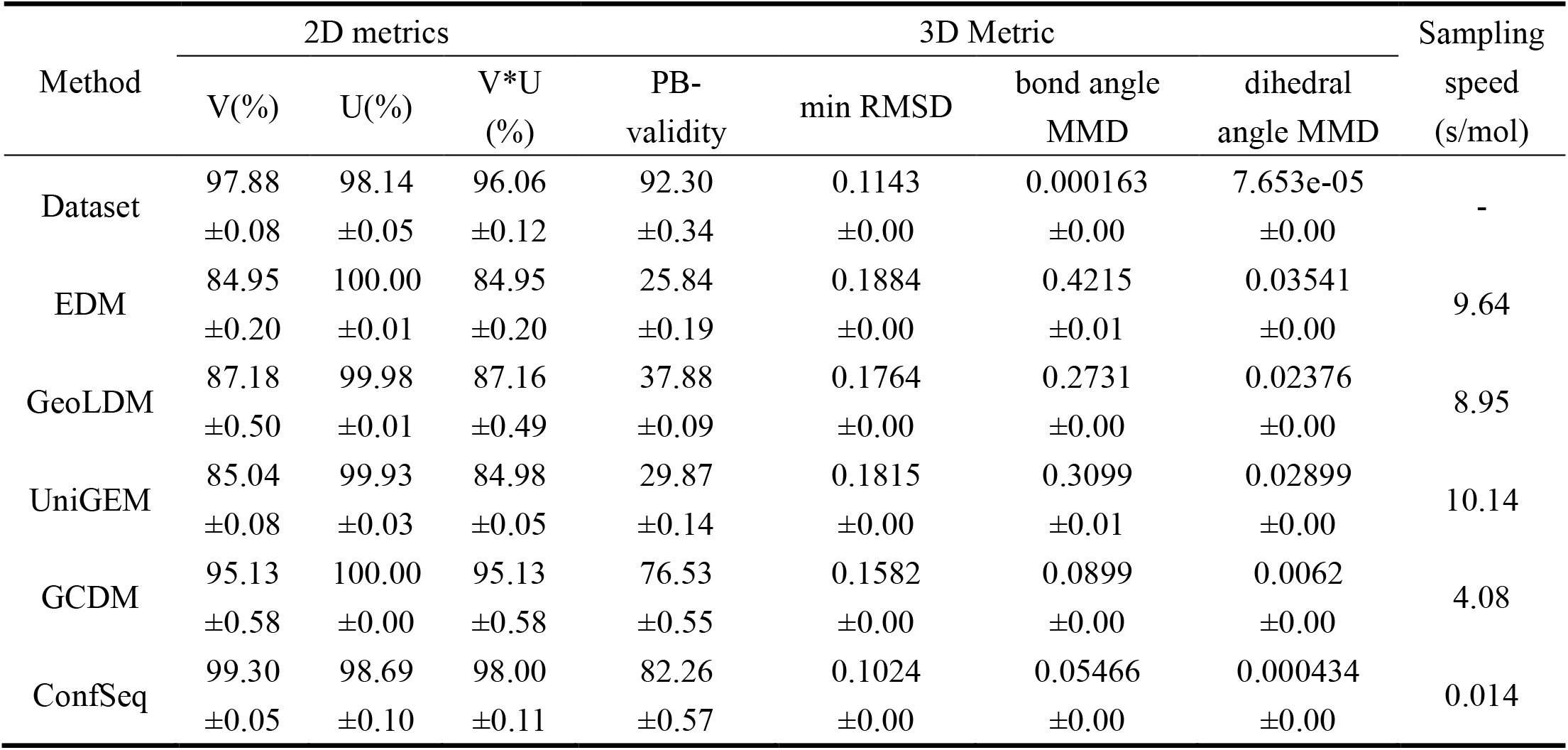
Performance comparison of 3D molecular generation methods in terms of 2D/3D metrics and sampling efficiency.

To further evaluate the similarity between the generated conformations and those in the training set, we computed the Maximum Mean Discrepancy (MMD) between the bond angle and dihedral angle distributions of the generated molecules and those of the reference dataset across 7 commonly occurring bond types. Compared to the previous state-of-the-art method, our model reduced the dihedral angle MMD from 0.0062 to 0.0004 and the bond angle MMD from 0.0899 to 0.0546. These results demonstrate that our model more accurately captures the conformational distribution of the training data than existing approaches. Notably, benefiting from the high inference efficiency of language models, our model achieves a sampling speed approximately 500 times faster than other diffusion-based methods.

Finally, we visualized the distribution of physicochemical properties and molecular representations of the molecules generated by different models. As shown in Fig. 3a and Supplementary Table 2, molecules generated by baseline methods exhibit substantial deviations from the training set in terms of QED, SAS, LogP, and TPSA. In contrast, the molecules generated by our model closely match the distribution of the training data across these properties. This observation is further supported by ChemNet embeddings(Fig. 3b)^32^, which show higher similarity between our generated molecules and those from the training set. Moreover, as illustrated in Fig. 3f, the 3D molecular fingerprints (E3FP) of our generated molecules also exhibit the highest resemblance to the training distribution (Fig. 3c)^33^. Together, these results demonstrate that our method produces molecules with the highest validity in both two-dimensional and three-dimensional evaluations. More importantly, the generated molecules maintain a high degree of consistency with the training set in terms of physicochemical properties and conformational distributions, which is unattainable for existing 3D molecular generation methods.

**Fig. 3.**
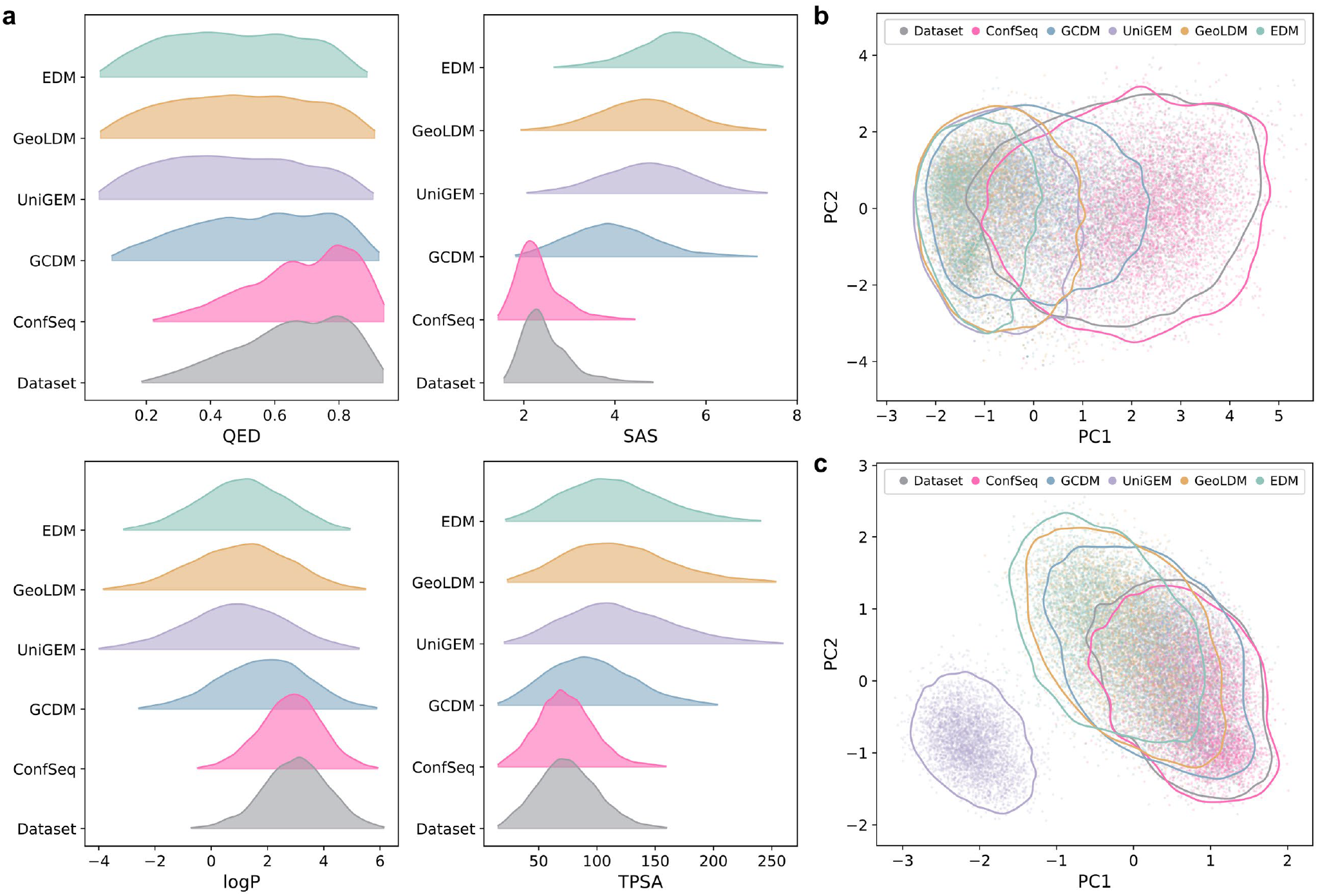
Comparison of the distributions of molecules generated by different methods. (a) Distributions of physicochemical properties, including QED, SAS, LogP, and TPSA. (b) PCA visualization of ChemNet embeddings of the generated molecules. (c) PCA visualization of E3FP molecular fingerprints of the generated molecules. The closed curves represent the distribution contours.

### Shape-conditioned 3D Molecule Generation with ConfSeq

Shape-conditioned 3D molecule generation is a long-standing need in drug design, enabling the creation of novel molecules that adopt spatial conformations similar to known active compounds while differing in their 2D structures. This strategy is particularly valuable for exploring novel chemical scaffolds and overcoming patent barriers.

To achieve shape-conditioned 3D molecule generation, we represent molecular surfaces as point clouds and use Rotation-Invariant Surface Convolution (RISConv) operations to extract invariant surface features^34^. These features are then fed into a Transformer decoder, which generates ConfSeq sequences that can be directly transformed into corresponding 3D molecular structures (Fig. 4a). We evaluated our method’s performance against existing approaches like ShapeMol and SQUID^35,36^. As shown in Fig. 4c, our method achieves validity comparable to ShapeMol and SQUID, with all methods nearing 100%. Crucially, our method demonstrates a significantly higher PB-validity score of 93.1%, surpassing ShapeMol (85.1%) and SQUID (70.1%). This superior PB-validity aligns with results from unconditional molecular generation tasks, further highlighting our model’s advantage in generating valid 3D molecules.

**Fig. 4.**
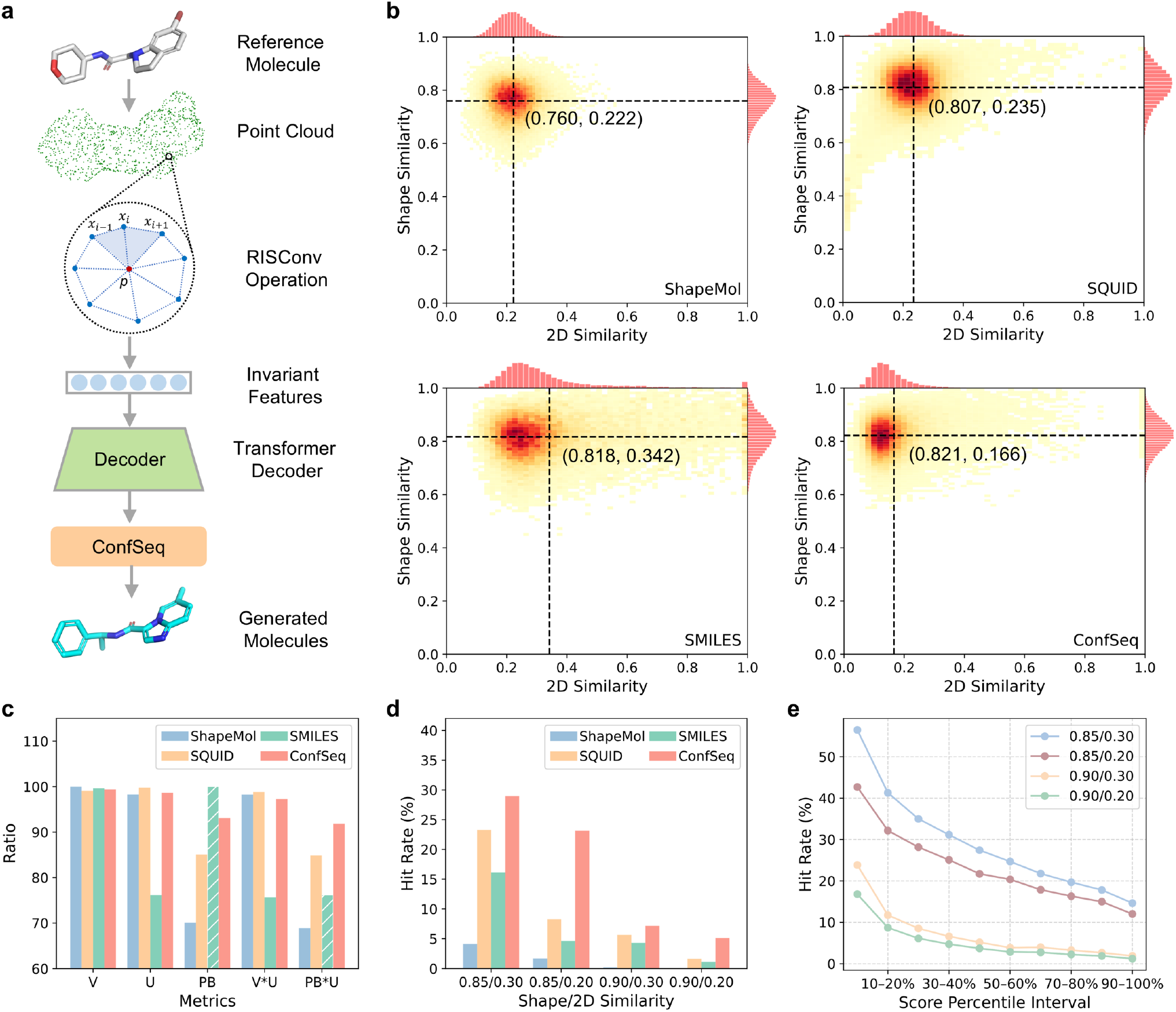
Comparison of different shape-conditioned 3D molecular generation methods. (a) Schematic illustration of the ConfSeq-based mode. (b) Histograms of shape similarity versus 2D similarity for molecules generated by different methods. The black line indicates the mean value. (c) Comparison of Validity (V), Uniqueness (U), PoseBuster validity (PB), and their product across different molecular generation methods. Bars with slashes indicate that conformers were generated using RDKit. (d) Hit rates of different models under various shape and 2D similarity thresholds. (e) Hit rates of molecules generated by the ConfSeq-based model, grouped according to their confidence score intervals.

We further compared the generated molecules with reference molecules in terms of shape and 2D similarity. As shown in Fig. 4b, molecules generated by our model exhibit higher shape similarity and lower 2D similarity on average. Using shape similarity thresholds (>0.85 or >0.9) and 2D similarity thresholds (>0.2 or >0.3) to define successful generations, our model achieved hit rates 24% to 210% higher than those of the best previous methods (Fig. 4e). Additionally, leveraging the intrinsic scoring capability of our language model, hit rates can be further improved through score-based filtering. For instance, under the thresholds of shape similarity >0.9 and 2D similarity <0.3, selecting the top 10% of generated molecules based on model-assigned scores increases the hit rate from 7.2% to 23.8%.

In previous studies, SMILES-based generation has been employed to design ligands with shapes complementary to protein pockets and host molecules^37,38^. In this work, we also implemented a SMILES-based method, where molecular conformations were reconstructed using RDKit. As shown in Fig. 4b–d, the SMILES-based method exhibited significantly inferior performance compared to our ConfSeq-based approach across key metrics, including molecular uniqueness, shape similarity, and 2D similarity. This comparison further highlights the superiority of ConfSeq over traditional SMILES-based methods in 3D molecular design.

### 3D Molecular Representation Learning with ConfSeq

Beyond 3D molecular generation, we further extended ConfSeq to 3D molecular representation learning. Specifically, we employed an encoder-only transformer to extract embeddings from the ConfSeq sequences of molecules, serving as vector representations of molecules. These representations were learned using a metric learning approach, wherein the distance between vectors reflects the 3D similarity of the corresponding molecules. This model is trained on 2.8 million molecules sourced from ChEMBL^39^ and BindingDB^40^, with conformations generated using RDKit.

We first applied the model to extract representations of ligands from the PDB database and visualized them by projecting into 2D using PCA^41^. As shown in Fig. 5a, the resulting representations successfully cluster some compounds that bind to the same protein pocket, even when their 2D structures differ substantially. For instance, the model captures the similarity between the binding conformations of linear inhibitors (2XFI_XFI and 4GID_0GH) and a macrocyclic inhibitor (4DPF_0LG) of the BACE-1 protein. This demonstrates the model’s ability to capture molecular 3D similarity.

**Fig. 5.**
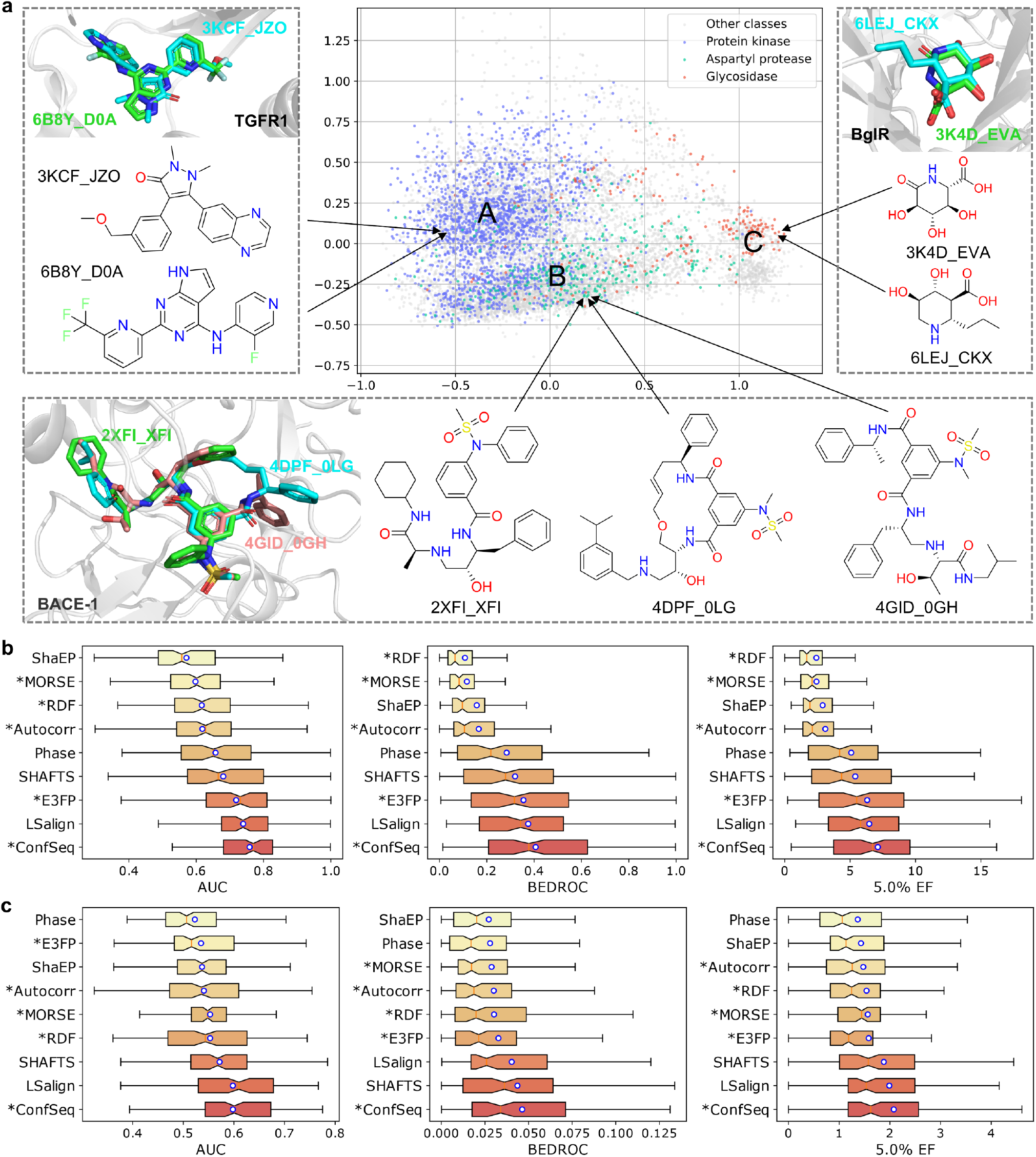
Performance of ConfSeq in 3D molecular representation learning. (a) PCA visualization of ligands from the PDB database using the proposed ConfSeq-based molecular representations. (b) Virtual screening performance of our representation and other methods on the DUDE dataset. (c) Virtual screening performance of our representation and other methods on the PCBA dataset. Blue circles represent the mean values. Models are ranked from top to bottom in ascending order of performance.

Additionally, we observed that the ligands in the PDB form three distinct clusters, with ligands targeting some proteins being enriched in particular clusters (Supplementary Fig. 4). For example, inhibitors of kinases, aspartyl proteases, and glycosidases predominantly reside in clusters A, B, and C, respectively. Further analysis of the molecules in each cluster revealed that cluster C predominantly contains small, highly polar molecules such as sugars, which explains the enrichment of glycosidase inhibitors in this cluster. cluster A is dominated by molecules with strong aromaticity and a high proportion of heterocycles, which is consistent with the typical features of kinase inhibitors. In contrast, molecules in cluster B exhibit relatively greater flexibility, contain more non-ring atoms, and possess a higher number of amide bonds. These characteristics suggest that molecules in cluster B are more likely to resemble certain peptides or protein fragments, which may account for the enrichment of some protease inhibitors in this cluster (Supplementary Fig. 5). These results indicate that our representation can capture some shared three-dimensional structural patterns among ligands targeting the same protein family, which are beyond the reach of existing 2D molecular representations (Supplementary Fig. 6).

In addition to visual analyses, we quantitatively evaluated the performance of our representation in ligand-based virtual screening using two well-established benchmark datasets, DUDE^42^ and PCBA^43^. As shown in the Fig 5b and 5c, our model achieves an AUC of 0.76, a BEDROC of 0.41, and a 5.0% enrichment factor (EF) of 7.12 on the DUDE dataset. For the PCBA dataset, the corresponding AUC, BEDROC, and 5.0% EF are 0.60, 0.046, and 2.08, respectively. It outperforms not only 3D molecular fingerprints such as E3FP^33^ and MORSE^44^, but also 3D similarity computation tools like LSalign^45^ and SHAFTS^46^. These tools rely on pairwise molecular alignment and similarity calculations, which are computationally intensive and time-consuming. In contrast, our approach computes molecular similarity directly via Euclidean distance between vector representations, achieving superior performance while delivering speedups by several orders of magnitude.

We further conducted virtual screening experiments using multiple query molecules, as shown in Supplementary Fig. 7. As the number of reference molecules increased, the screening performance of the model also improved. This suggests that, with sufficient reference molecules, the efficiency advantage of our method can be further translated into performance gains. To facilitate large-scale applications, we have precomputed molecular embeddings for all three-dimensional compounds in PubChem using our model. Based on these representations, the 3D similarity between a query molecule and the entire set of 98 million PubChem compounds can be computed within one minute on a single CPU core, enabling ultra-fast, large-scale virtual screening.

## Conclusion

In this work, we addressed the significant challenge of extending the success of CLMs from 2D to 3D molecular modeling, which has been hindered by the lack of effective 1D representations for 3D molecular structures. We introduced ConfSeq, a novel molecular conformation description language that integrates SMILES notation with internal coordinates. This approach inherently ensures SE(3)-invariance while maintaining the desirable properties of SMILES, such as readability and conciseness.

ConfSeq enables the reformulation of diverse 3D molecular modeling tasks, such as conformation prediction, unconditional and shape-conditioned 3D molecular generation, and 3D molecular representation, into sequence modeling problems amenable to standard language model architectures like the Transformer. Our results demonstrate that this approach achieves state-of-the-art performance across various benchmark datasets, often significantly outperforming established methods. Several distinctive advantages of the ConfSeq-based approach over prevailing diffusion-based methods in 3D molecular modeling were highlighted throughout our study. These include markedly improved inference efficiency, achieving molecular generation sampling speeds approximately 500 times faster; the ability to control the diversity of generated structures through sampling temperature; and the capability to score and rank generated candidates. This scoring capability enables efficient selection of high-quality molecular candidates.

In summary, ConfSeq provides an efficient bridge between the power of sequence-based language models and the complexities of 3D molecular structures. We believe ConfSeq serves as a foundational tool that can catalyze the development of CLMs, potentially establishing a new and promising paradigm for AI-driven 3D molecular modeling in chemistry and drug discovery.

## Method

### Details of ConfSeq

ConfSeq encodes a series of custom symbols to represent the conformational information of molecules. The extraction of conformational information and the generation of ConfSeq are jointly performed using two chemical informatics packages, RDKit and Indigo. Three types of conformational information are included, as follows:

The first type is the dihedral angle, defined by four consecutively bonded atoms (i–j–k–l). It describes the angle between the plane formed by atoms i–j–k and the plane formed by atoms j–k–l. Dihedral angles are the most critical determinants of molecular conformation. For each rotatable bond (j–k), multiple dihedral angles may arise due to different choices of adjacent atoms i and l, as well as the orientation of the central bond, resulting in distinct dihedral pathway. To address this, we implement a standardization algorithm that assigns a unique dihedral pathway to each rotatable bond. The direction of the central bond is determined by ordering the two atoms so that the one with the smaller index comes first. When selecting the neighboring atoms i and l, priority is given to atoms with a degree greater than one. If multiple atoms satisfy this condition or none do, the atom with the smallest index is selected. Subsequently, a unique dihedral angle φ is then calculated for each rotatable bond, rounded to the nearest integer, and encapsulated in angle brackets to form a discrete token, such as “<113>”. This token replaces the corresponding bond tokens in the explicit-bond SMILES representation. In previous studies, dihedral angles have typically been considered only for rotatable bonds. In this work, we additionally incorporate dihedral angles within non-aromatic rings, which also play a crucial role in defining molecular conformations.

The second type is the bond angle, which is defined by three consecutively bonded atoms (i–j–k) and measures the angle formed at the central atom (j) between the two bonds connecting it to atoms i and k. In ConfSeq, bond angles are considered where the central atom is located outside of any ring and bonded to two heavy atoms (excluding oxygen atoms bonded via double bonds). Compared to central atoms that are constrained within rigid ring systems or bonded to multiple neighboring atoms, these bond angles exhibit greater intrinsic flexibility and have a more pronounced influence on the overall molecular conformation. Similar to the torsion angle, the bond angle value is rounded and enclosed in angle brackets, followed by a pipe symbol (“|”) to distinguish it from torsion angle tokens. Each bond angle token (e.g., “<30> |”) is then inserted into the ConfSeq sequence immediately after the token representing the central atom (j).

The third type is pseudo-chirality. For a rotatable bond formed between two chiral centers, a single dihedral angleis sufficient to define the relative spatial arrangement of the two central atoms and their neighboring groups.

However, in molecules containing non-chiral atoms, as illustrated in Supplementary Fig. 8, the dihedral angle 1–2–5–7 can only determine the relative positions of atoms 1, 2, 3, 4, 5, and 7. The position of atom 6 remains ambiguous because it can adopt two interchangeable configurations. These two configurations can rapidly interconvert through lone pair inversion and are chemically indistinguishable, hence, the nitrogen atom is not considered chiral. Nevertheless, in 3D molecular modeling, it is necessary to distinguish between these two configurations. To address this, we introduce the concept of pseudo-chirality to differentiate them. Specifically, the symbols “}” and “{” are used to denote counterclockwise and clockwise pseudo-chiral centers, respectively. These symbols are placed before or after the corresponding dihedral angle token, depending on the relative atomic indices of the pseudo-chiral atoms.

### Dataset

Our study involved multiple computational experiments. For conformation prediction, we used a subset of the GEOM-Drugs dataset curated by Zhang et al., following their predefined data splits^47^. The training and validation sets contain 39,990 and 5,000 molecules respectively, each with 5 conformers per molecule. The test set includes 200 molecules and 14,317 conformers in total. Unconditional 3D molecular generation experiments were also conducted on another subset of the GEOM-Drugs dataset, as introduced by Hoogeboom et al^48^. This subset contains 430,000 molecules, for each of which the 30 lowest-energy conformers were selected. The dataset was partitioned into training, validation, and test sets following an 8:1:1 ratio. The molecular structures, initially provided in XYZ format, were first converted into SDF format using OpenBabel for subsequent analysis. For shape-conditioned 3D molecular generation, we followed the experimental protocol established by Chen et al.^36^, utilizing the MOSES dataset along with their RDKit-generated single 3D conformers for each molecule. We adhered to their predefined data splits, which included 1,592,653 molecules for training, 1,000 for validation, and 1,000 for testing. For molecular representation learning, we constructed a dataset of 2,769,534 molecules sourced from ChEMBL^39^ and BindingDB^40^, generating 10 conformers per molecule using RDKit. We then created a training dataset consisting of 95.7 million randomly sampled molecular pairs, with similarities calculated using LSalign^45^. The protein-ligand complex crystal structure data were sourced from Wang et al.^49^. Ligands were deduplicated based on the extended-connectivity fingerprints (ECFP) similarity, using a threshold of 0.7. Additionally, protein category information was retrieved from the UniProt database via its official API^50^. We performed ligand-based virtual screening using the DUD-E and PCBA datasets, adhering to the evaluation protocol outlined by Jiang et al.^51^. The DUD-E dataset comprises 102 pharmacological targets, with an average of 224 active ligands per target. For each active ligand, approximately 50 decoy compounds were generated. The PCBA dataset includes 15 target sets, consisting of 7,761 active ligands and 382,674 unique inactive compounds, sourced from the PubChem Bioassay data.

### Implementation of the ConfSeq-based model

For the conformation prediction task, we employed a Transformer-based architecture consisting of 6 encoder layers and 6 decoder layers, with a model dimension of 256. The model’s input is a modified SMILES sequence, generated by inserting placeholder tokens at specific positions within the original SMILES string to ensure alignment with the corresponding ConfSeq sequence. To enhance the diversity of the training data, we employed 100-fold data augmentation. This approach involved generating multiple distinct pairs of input sequences and ConfSeq sequences for each 3D molecule, utilizing a strategy similar to SMILES enumeration. Similarly, during inference, we randomly sampled and generated multiple different input sequences for a given molecule.

For the unconditional molecular generation task, we adopted a decoder-only Transformer architecture consisting of 6 decoder layers and a hidden dimension of 768. We applied 5-fold data augmentation during training to further improve model performance. For the shape-conditioned generation task, we followed the approach of Chen et al.^36^, first constructing the molecular surface mesh for each molecule, then uniformly sampling 1,024 points on the mesh with probabilities proportional to the corresponding face areas. From these sampled points, we derived highly expressive, rotation-invariant surface descriptors. These descriptors were processed using RISurConv, an attention-augmented convolutional operator that incorporates self-attention mechanisms to enhance the representational capacity of surface features. The resulting invariant features were subsequently fed into a Transformer decoder to guide the generation of 3D molecular structure. This decoder shares the same architecture as that used for the unconditional generation task, comprising 6 layers with a hidden dimension of 768. Across all aforementioned tasks, the models were trained using the cross-entropy loss function. During inference, ConfSeq sequences were generated using a random sampling strategy. The generation score for each predicted sequence was calculated as the average log-probability across all tokens.

For the molecular representation task, we employed an encoder-only Transformer architecture consisting of 6 layers with a hidden dimension of 256. A Siamese network framework was adopted to extract representations for pairs of molecules. The similarity between two molecular embeddings, *s*(*q, f*), was computed based on their Euclidean distance *d*(*q, f*), defined as:

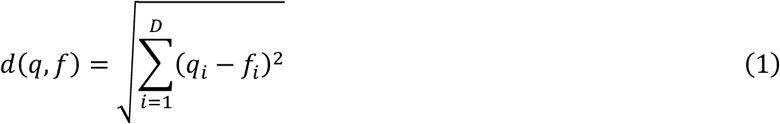

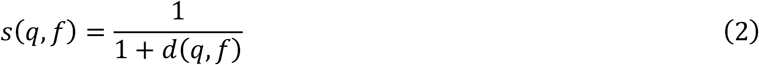

where *q, f* denote the embeddings of the query molecule and the reference molecule, respectively, and *D* is the embedding dimension.

### Evaluation metrics and baseline methods

In the conformation prediction task, we compared our method against SEDGEN^47^, GeoDiff^52^, UniMol^53^, DMCG^54^, GADIFF^55^, and Torsional Diffusion (Tor. Diff.)^20^, EC-Conf^56^, CGCF^57^, ConfVAE^58^, CONFGF^59^, RMCF^60^, GeoMol^61^, DGSM^62^, AGDiff^63^, Tora3D^64^. The results for Tor. Diff. and DMCG were obtained from our own training, while the results for SEDGEN, GeoDiff, and UniMol were derived from author-released checkpoints. The results for the remaining models were directly cited from prior reports.

We assess the diversity and quality of generated molecular conformers using Coverage (COV) and Matching (MAT) scores. The COV score quantifies the proportion of structures in one set that are covered by another set, while the MAT score calculates the average RMSD between each structure in one set and its closest counterpart in the other set. COV and MAT metrics, following the conventional recall-based evaluation (COV-R, MAT-R), can be defined as follows:

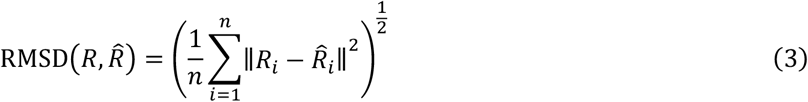

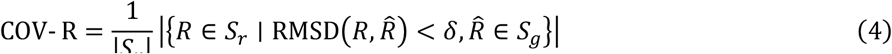

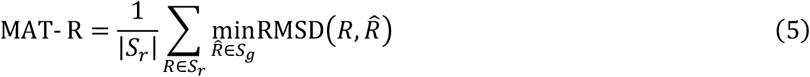

where the threshold δ for the Coverage score is set to 0.75 Å, *S*_*g*_ denoting the set of generated conformers and *S*_*r*_ denoting the set of reference conformers for a given molecule, *S*_*g*_ is set to be twice the size of *S*_*r*_ for each molecule. COV-P and MAT-P are defined analogously, with the generated and reference conformer sets exchanged in the respective formulations.

For unconditional molecular generation, we followed prior work and conducted three independent rounds of random sampling per model, generating 10,000 molecules in each round. For the benchmark models (EDM^48^, GeoLDM^65^, GCDM^66^, and UniGEM^67^), we utilized official checkpoints and converted the generated XYZ files into SDF format using OpenBabel before evaluation. We evaluated the generated molecules across various metrics, including validity, uniqueness, and PB-validity. Molecular properties, including the Quantitative Estimate of Drug-likeness (QED), Synthetic Accessibility Score (SAS), logarithm of the octanol–water partition coefficient (LogP), and Topological Polar Surface Area (TPSA), were calculated using RDKit. Additionally, ChemNet activation embeddings were extracted using the fcd_torch package^32^, and E3FP fingerprints were computed using scikit-fingerprints^33^.

For the shape-conditioned generation task, we sampled 50 ConfSeq sequences for each reference molecule. As baselines, we directly employed the generated molecules provided in the studies of SQUID^35^ and ShapeMol^48^, which used the same reference molecules as ours. Similar to the unconditional generation setting, we evaluated the validity, uniqueness, and PB-validity of the generated molecules. Shape similarity scores were computed using the freely available ShaEP software^68^, configured to depend solely on molecular shape features. The ECFP fingerprints and Tanimoto similarity were computed using RDKit to quantify 2D similarity. Finally, we reported the mean shape similarity and mean 2D similarity between generated molecules and their corresponding references.

For molecular representation evaluation, we compared our method against 3D similarity computation tools (ShaEP^68^, SHAFTS^46^, LSalign^45^, and Phase^69^) as well as 3D molecular fingerprints (RDF, MORSE, Autocorr and E3FP) the latter computed using Scikit-fingerprints^70^. The evaluation was conducted using the area under the ROC curve (AUC), the Boltzmann-enhanced discrimination of ROC (BEDROC) at α = 80.5, and the enrichment factor at the top 5.0% (5.0% EF).

## Acknowledgments

We gratefully acknowledge financial support from National Natural Science Foundation of China (grant no. T2225002 and 82273855, M.Z.), National Key Research and Development Program of China (grant no. 2022YFC3400504 and 2023YFC2305904, M.Z.), the Strategic Priority Research Program of the Chinese Academy of sciences (grant no. XDB0830200, M.Z.), the open fund of state key laboratory of Pharmaceutical Biotechnology, Nanjing University, China (grant no. KF-202301, M.Z.), and Shanghai Post-doctoral Excellence Program (grant no. 2024707, J.X.).

## Author Contributions

J. Xiong proposed the idea, and conducted computational experiments and drafted the initial manuscript together with Y. Shi. W. Zhang and R. Zhang participated in the analysis of results. Z. Chen, C. Zeng and X. Jiang contributed to the case analysis. W. Zhang, D. Cao, Z. Xiong, and M. Zheng helped check and improve the manuscript. M. Zheng led the project and designed the study. All authors read and approved the final manuscript.

## Competing Interests

The authors declare that they have no competing interests.

## Notes

### Competing Interest Statement

The authors have declared no competing interest.

