## Supplementary Information for "Bridging 3D Molecular Structures and Artificial Intelligence by a Conformation Description Language"

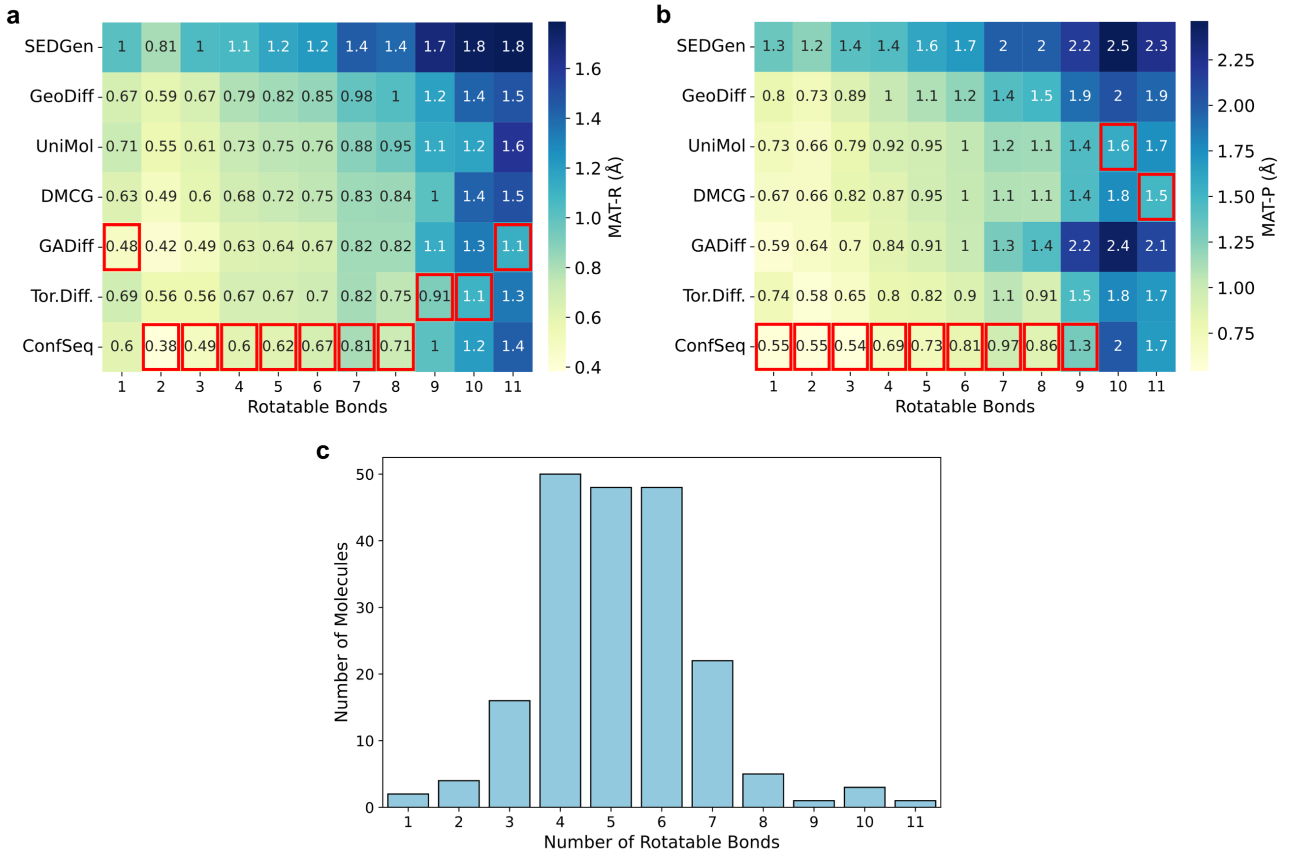


Supplementary Fig. 1. Performance comparison of different models on molecules with varying numbers of rotatable bonds. (a–b) MAT-R and MAT-P values for each model, respectively, with the best values highlighted in red boxes. (c) Distribution of molecules based on the number of rotatable bonds.

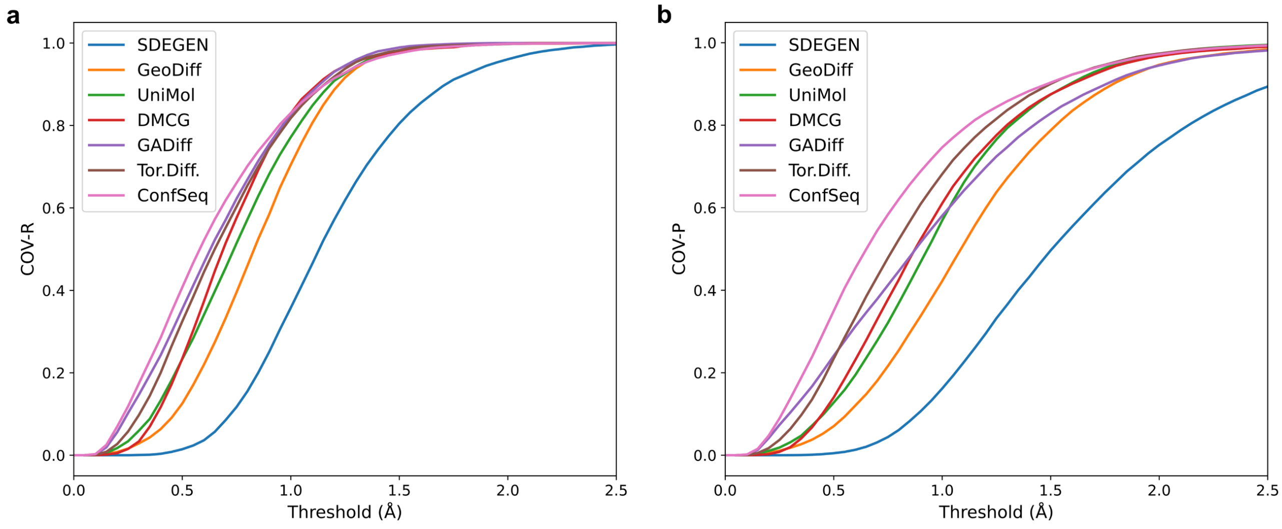


Supplementary Fig. 2. COV-R (a) and COR-P (b) scores for different model at various threshold levels.

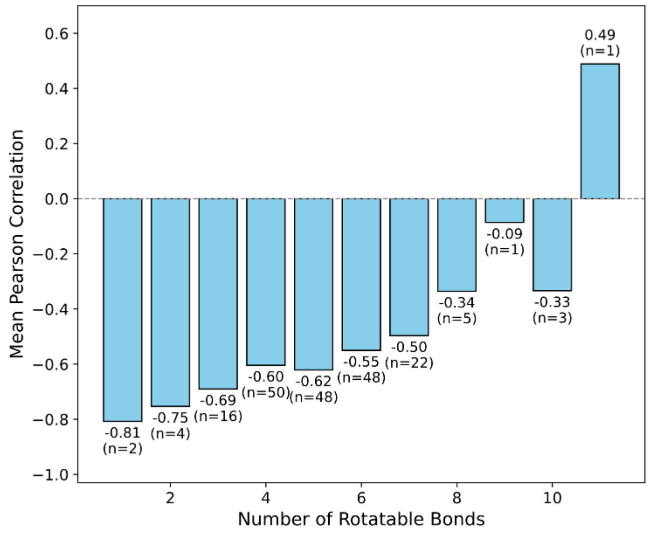


Supplementary Fig. 3. Pearson correlation coefficient between the energy and scores for molecules with varying numbers of rotatable bonds.

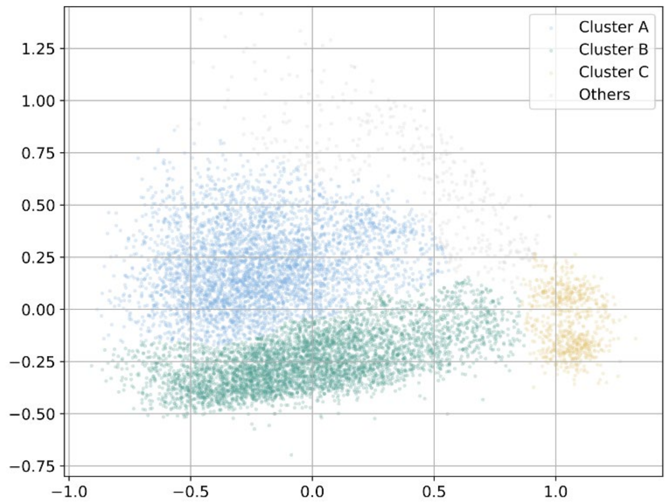


Supplementary Fig. 4. Clustering of PDB ligands based on ConfSeq-derived molecular representations using a Gaussian mixture model.

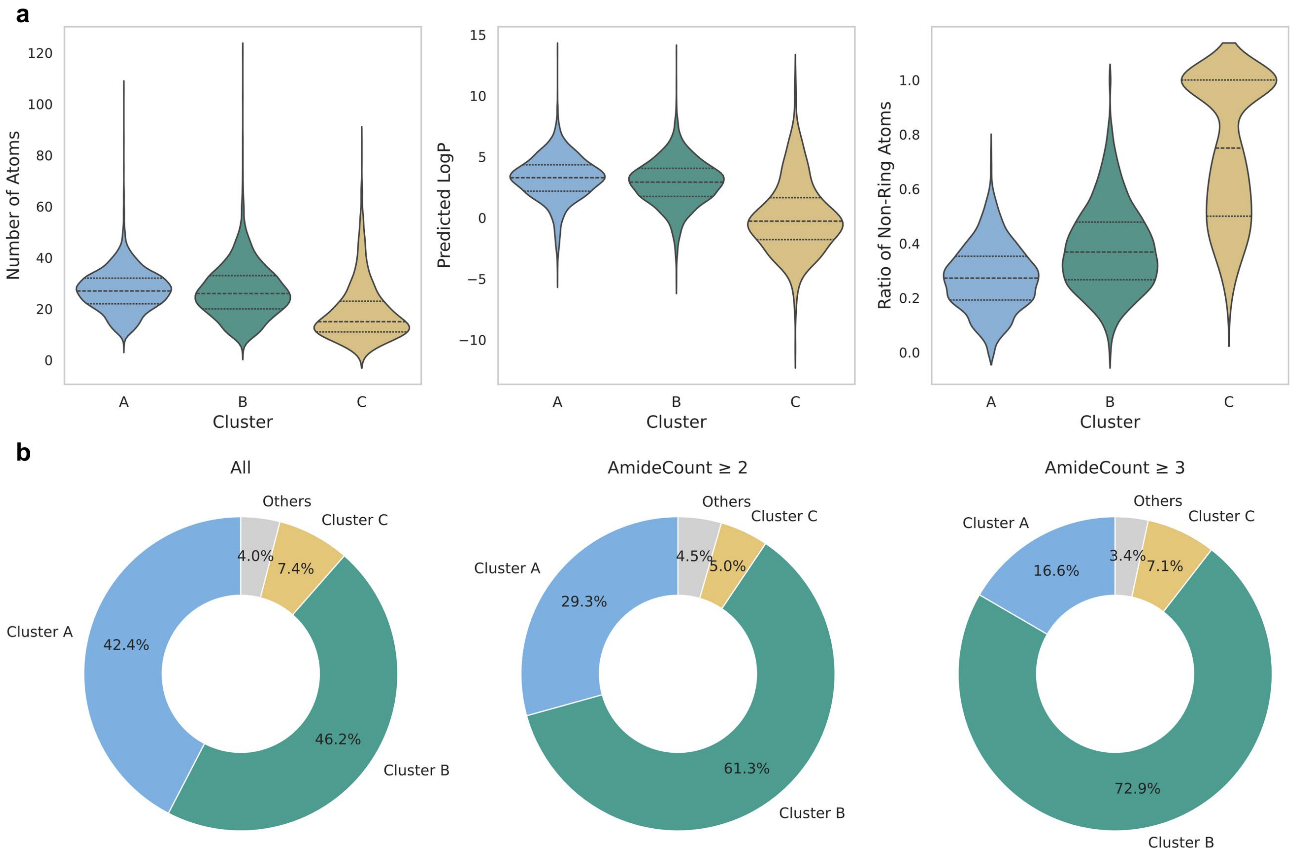


Supplementary Fig. 5. (a) Comparison of the distributions of atom count, LogP, and the ratio of non-ring atoms across Clusters A, B, and C. (b) Proportional distribution of molecules assigned to Clusters A, B, C, and Others for the full dataset (All), as well as for subsets containing molecules with ⩾2 and ⩾3 amide bonds.

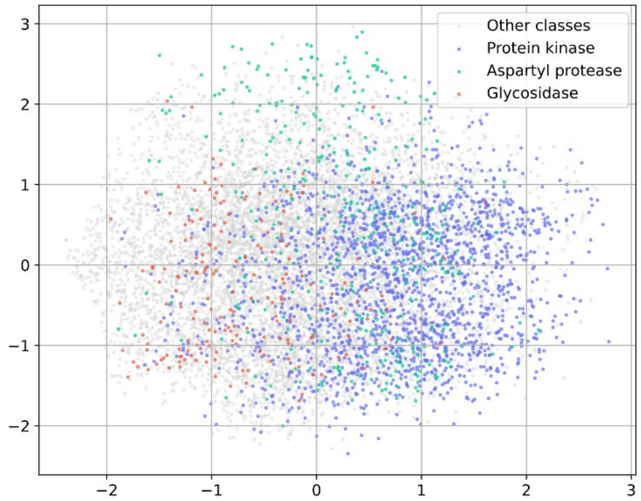


Supplementary Fig. 6. PCA visualization of ligands from the PDB database using ECFP molecular fingerprints.

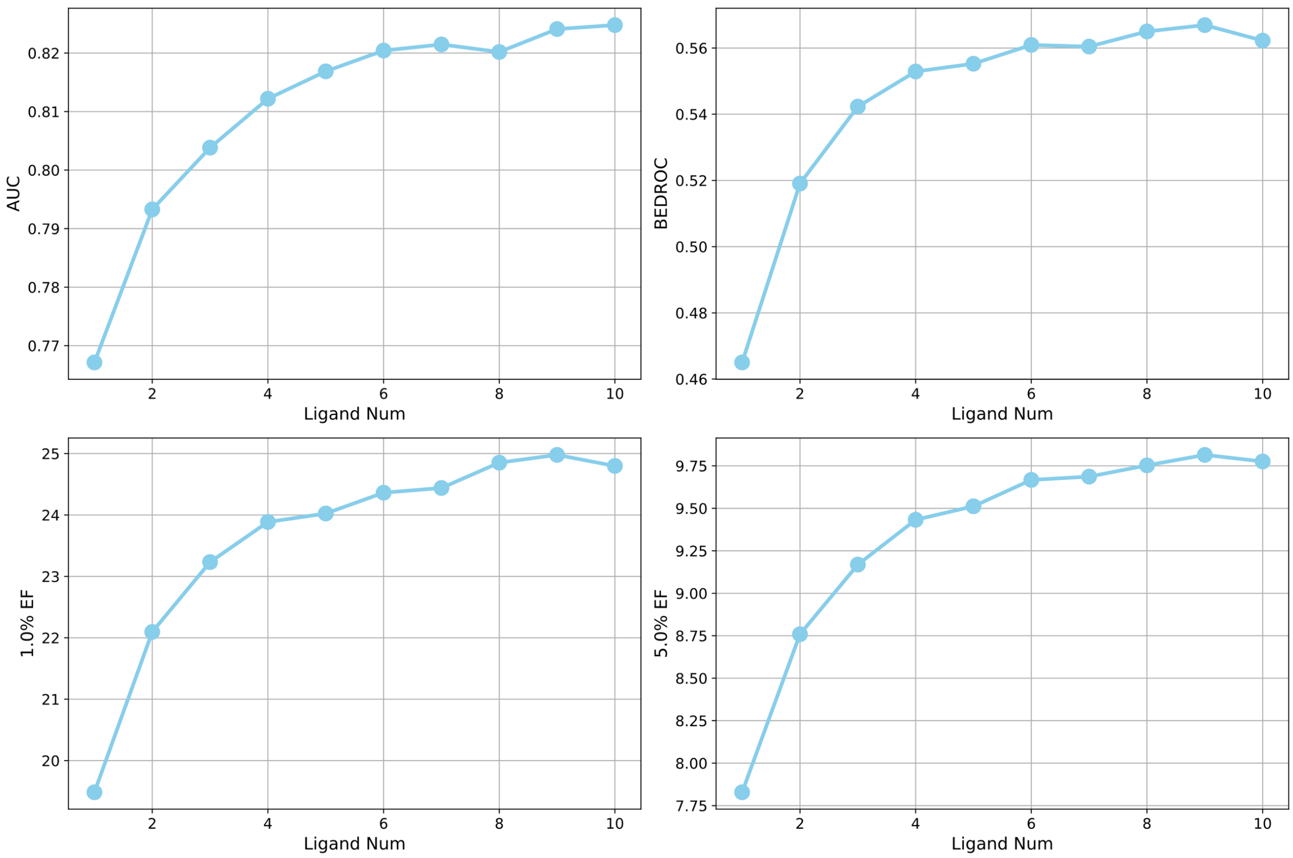


Supplementary Fig. 7. Impact of the number of reference ligands on virtual screening performance, measured by AUC, BEDROC, and enrichment factors at 1.0% and 5.0%.

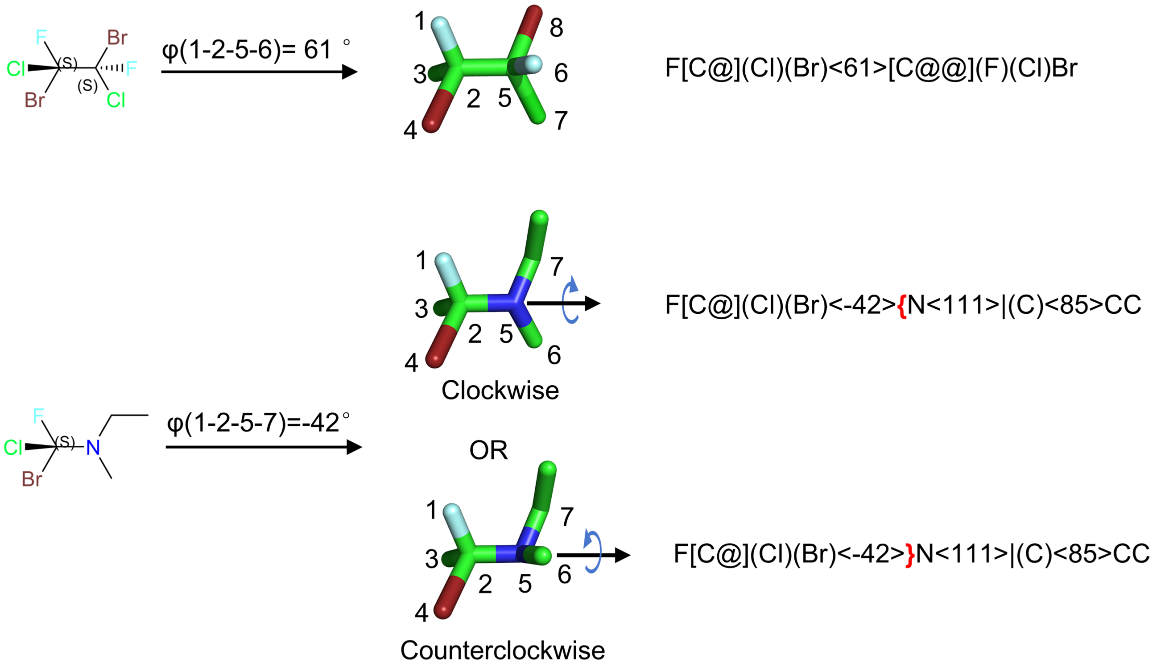


Supplementary Fig. 8. Schematic illustration of the proposed pseudo-chirality concept.

Supplementary Table 1. Performance of different models on conformation prediction for the GEOM-DRUG dataset

| Models | COV-R(%)↑ | | MAT-R(Å)↓ | | COV-P(%)↑ | | MAT-P(Å)↓ | |
|---|---|---|---|---|---|---|---|---|
| | Mean | Median | Mean | Median | Mean | Median | Mean | Median |
| SEDGen | 11.70 | 1.79 | 1.1840 | 1.1528 | 4.42 | 0.71 | 1.6238 | 1.5818 |
| GeoDiff | 39.54 | 33.75 | 0.8405 | 0.8255 | 21.66 | 15.39 | 1.1443 | 1.1103 |
| UniMol | 51.43 | 50.42 | 0.7625 | 0.7531 | 32.14 | 32.14 | 0.9838 | 0.9761 |
| DMCG | 57.60 | 62.75 | 0.7331 | 0.6998 | 37.82 | 35.98 | 0.9623 | 0.9137 |
| GADIFF | 62.23 | 63.61 | 0.6674 | 0.6491 | 41.27 | 37.02 | 0.9800 | 0.9121 |
| Tor.Diff. | 60.29 | 64.91 | 0.6963 | 0.6674 | 47.88 | 47.73 | 0.8638 | 0.8293 |
| ConfSeq | **66.05** | **69.05** | **0.6488** | **0.6295** | **58.38** | **59.05** | **0.7738** | **0.7204** |

Note: ↑ indicates that higher values are better, ↓ indicates that lower values are better, and the best results are highlighted in bold.

Supplementary Table 2. Distributional differences of molecular properties between generated and training set molecules

| Models | QED | SAS | LogP | TPSA |
|---|---|---|---|---|
| EDM | 0.181152 | 2.854648 | 1.914085 | 38.563136 |
| GeoLDM | 0.160342 | 2.140837 | 1.966582 | 42.228730 |
| UniGEM | 0.182138 | 2.222610 | 2.125428 | 43.924866 |
| GCDM | 0.101732 | 1.461064 | 1.168936 | 19.939621 |
| ConfSeq | 0.028951 | 0.163774 | 0.181050 | 2.122042 |

Note: The distributions are compared using the Wasserstein distance computed with the SciPy library. Lower values indicate greater similarity to the training set distributions.
